# Genomic Correlates of Tailocin Sensitivity in *Pseudomonas syringae*

**DOI:** 10.1101/2023.04.24.538177

**Authors:** David A. Baltrus, Savannah Weaver, Laura Krings, Anh Evy Nguyen

**Affiliations:** School of Plant Sciences, University of Arizona, Tucson AZ, USA; School of Animal and Comparative Biomedical Sciences, University of Arizona, Tucson AZ, USA; Department of Molecular and Cellular Biology, University of Arizona, Tucson AZ, USA

## Abstract

Phage derived bacteriocins, also referred to as tailocins, are structures encoded by bacterial genomes and deployed into the extracellular environment to target and kill sensitive cells. Tailocins display great potential as agricultural antimicrobials due to their durability, efficiency, and specificity of killing with prophylactic application of these molecules having been shown to prevent infection by multiple phytopathogens. Although previous reports have strongly suggested that tailocins of *Pseudomonas syringae* bind sugar moieties in the LipoPolySaccharide (LPS) of target cells, the molecular mechanisms and binding interactions that enable tailocins from *P. syringae* to kill sensitive targets remain unclear. We therefore carried out a genome-wide association study investigating tailocin sensitivity across a diverse set of *P. syringae* genomes. Our results demonstrate that genes strongly correlated with tailocin sensitivity are localized to one contiguous region on the *P. syringae* chromosome encoding LPS structures similar to the Common Polysaccharide Antigen of *P. aeruginosa*. We further find that enzymes involved in the biosynthesis and transport of D-rhamnose and L-rhamnose are associated with tailocin sensitivity classes A and B, respectively, with large-scale recombination of the O-antigen biosynthesis region likely underlying rapid and fundamental changes in LPS structure between strains. We identify the *rfbD* gene as an additional genomic indicator to predict tailocin sensitivity and use this information to test tailocin interactions with unscreened strains from across phylogroups, including some in which LPS chains have been previously characterized. Overall, our results strongly support that tailocin sensitivity across *P. syringae* strains is largely determined by recombination events across strains that lead to differential production of either D or L-rhamnose moieties in the main LPS chain.

## Introduction

Phage derived bacteriocins, hereafter referred to as tailocins, are proteinaceous structures encoded by bacterial genomes thought to be used by strains in an extracellular manner to target and kill competing strains under natural conditions[1–3]. The efficacy of killing, durability, and strain specificity of tailocins have made them exceptional candidates for development and deployment as antimicrobials in clinical and agricultural environments[4, 5]. To best maximize the efficiency of use and develop methods to optimize application strategies, as well as to predict sensitivity of untested strains, we must better understand the genetic basis of resistance to tailocins across diverse suites of strains. Here, we present a genome wide association analysis of loci that affect tailocin based killing within *Pseudomonas syringae* strains across a set of phylogenetically diverse strains previously screened for tailocin sensitivity and use this information to predict tailocin sensitivity for additional strains from their genome sequences.

Tailocins are part of a broad class of structures, also including type VI secretion systems and extracellular contractile injection systems (eCIS), coopted by bacterial genomes from phages and structurally resembling phage tails[2, 3]. They are produced by bacterial cells and released through cell lysis, but unlike either the type VI systems or eCIS, are not currently known to deliver effector proteins to target cells. Tailocins have evolved at least multiple times independently across taxa from different phage: with the R-type tailocins evolving independently from various myoviridae phage and F-type tailocins are derived from a phage in the siphoviridae family [1, 6]. Although a variety of bacteria besides Pseudomonads have been shown to produce tailocins, much of our current knowledge concerning tailocin killing and resistance comes from studies investigating R-type tailocins (also known as R-type pyocins) of *P. aeruginosa* [2, 3, 7–11]. R-type pyocins have been shown to efficiently kill sensitive cells through direct lysis by mechanical disruption of target cell membranes, with binding of tailocins determined largely by receptor binding proteins (Rbps) and their associated chaperones[8, 9, 12]. These two loci together appear to determine the targeting spectrum for both types of Pseudomonad R-type tailocins (pyocins and syringocins), with the targeting spectrum for R-type syringacins found in *Pseudomonas syringae* has been shown to rapidly evolve through localized recombination of regions containing the Rbps [13].

The outer membranes of Gram negative bacteria are decorated with a variety of proteins and sugars, with one of the most well-known structures being the lipopolysaccharide (LPS) layer [14]. The LPS is composed of covalently bound chains of various sugar moieties, the compositions of which vary within and between species, anchored into the outer membrane [15]. Since the LPS is a major component determining interactions of bacteria with the immune systems of hosts, is often used for attachment of phage, and is likely involved in a variety of other processes important for survival across bacterial populations and communities, composition of the LPS is thought to be under strong diversifying selection and can be notoriously variable even between closely related strains [16]. Earlier studies on the R-pyocins of *P. aeruginosa* implicated the LPS as the binding sites for these tailocins and specifically by the Rbps, and pointed towards either rhamnose or glucose as the interacting sugar moieties [9]. Additionally, *P. aeruginosa* strains are able to produce multiple types of O-antigen that are potentially linked to the LPS core: one type of chain is referred to as the Common Polysaccharide Antigen (CPA) or A band and is composed of repetitive chains of D-rhamnose linked together, while the other chain can be composed of more diverse sugars and is referred to as the B band [18]. The presence, composition, and length of these different O-antigen bands has been previously implicated as playing a role in shielding or enabling attachment of tailocins to LPS-associated sugars [9, 18]. Further, RB-TnSeq across a handful of *P. fluorescens* strains implicated both the core and O-antigen bands of the LPS as interaction sites for these molecules, with disruption of genes from all of these regions providing resistance against tailocin-based killing [19]. Conversely, multiple papers have implicated the Common Polysaccharide Antigen or A band of the LPS as a main site of interactions of tailocins from *P. syringae* as genetic data suggests that *P. syringae* strains may only produce variations on the CPA and deletions or modification of genes within the CPA production pathway differentially affect tailocin targeting to otherwise sensitive strains [20, 21]. Indeed, in natural populations of *P. viridiflava* (classified within the *P. syringae* species complex and sharing vertically inherited tailocin structural loci as *P. syringae*) LPS conformation is correlated with tailocin sensitivity [22]. This difference in tailocin targeting between R-pyocins and R-syringacins is perhaps not completely surprising as it echoes from our current understanding that tailocins of *P. syringae* are derived from a different progenitor phage than the R-type pyocins of *P. aeruginosa* and many of the R-type tailocins of *P. fluorescens* [6].

Here, we report a genome-wide association study to map the genetic basis of tailocin sensitivity across a variety of closely related *P. syringae* strains and in order to identify genomic regions associated with differing sensitivity to R-type syringacins. We then extrapolate from these results to map and predict tailocin sensitivity across a diverse range of isolates from this species, including numerous strains where LPS has been previously characterized, and uncover genomic trends that appear to enable prediction of tailocin sensitivity for these strains. Overall, our results highlight the variability of pathways determining LPS structures throughout *P. syringae*, suggest that large-scale recombination of one region on the chromosome is responsible for fundamental shifts in LPS composition in *P. syringae*, and provide information that could enable prediction of tailocin sensitivity across this phytopathogenic species. All evidence strongly suggests that *P. syringae* tailocins largely interact with either the L or D-rhamnose moieties of the LPS.

## Methods

### Genome Sequences Used for Genome-Association Studies and Tailocin Prediction Studies

All genomes referenced in this manuscript are listed in Table 1, which also lists accessions for these sequences at Genbank and abbreviations used for each strain in the manuscript. We additionally report draft and complete genomes for a variety of strains that have had their LPS conformations previously characterized in [23], as well as complete genome sequences for two closely related strains (CC1417 and CC1524) containing plasmids that house numerous genes predicted to affect LPS biosynthesis. Methods and statistics for genome sequencing and assembly for LPS-characterized strains as well as CC1417 and CC1524 are described in File S1 at doi: 10.6084/m9.figshare.22688020. Software used for genome assemblies can be found in File S1[24–27].

### Genome-wise Association with Tailocin Sensitivity

We downloaded .fasta sequences for each of the genomes in Table 1 that had been previously screened for tailocin sensitivity in [13], and used Prokka to reannotate each in a consistent manner [28]. We then carried out a clustering analysis of the .gff files from these genomes using Roary and with default parameters [29]. Last, we used Scoary to associate tailocin sensitivity with clusters of orthologous genes found by Roary [30]. Tailocin sensitivity classes for each gene as used in the Scoary analysis are listed in Table 1, and we note that sensitivity classes for strains CC1630, *Pto*T1, and *Pla*106 differ from what was published in [13] due to labelling errors in the previous manuscript. All input and output files for Prokka, Roary, and Scoary analyses can be found as supplemental files at doi: 10.6084/m9.figshare.22688020.

### Phylogenetic Analysis

We obtained protein sequences for RfbD from each of the genomes listed in Table 1, and aligned these using Clustal Omega with default parameters [31]. We used Modeltest to find the best evolutionary model describing this alignment [32], which was JTT+G4. We then inferred phylogenetic relationships from this alignment using RaxML-ng [33]. We used the following command to infer phylogenies and bootstrap support:

`raxml-ng --all -msa Alignment.fasta --model JTT+G4 --prefix Prefix -- bs-metric fbp,tbe`

. All input and output files for phylogenetic analyses can be found as supplemental files at doi: 10.6084/m9.figshare.22688020.

### Soft-agar Overlay of Previously Unscreened Strains for Tailocin Sensitivity

Soft-agar overlays were carried out as previously described [34]. Briefly, overnight cultures of strains CC440, USA011, USA007, and CC1416 were diluted back 1:100 in King’s B media and grown for 4 hours at 27°C while shaking at 220rpm. At this point, mitomycin C was added to these cultures to a final concentration of 0.75 µg/mL and the cultures were incubated under the same conditions overnight. The next day, tailocins from these strains were purified from the supernatant by filtration through 0.22 µm syringe filters.

For overlays, strains of interest were grown overnight in KB media at 27°C while shaking at 220rpm. The next morning, cultures were diluted 1:100 in fresh KB media and incubated an additional 4 hours under the same environmental conditions. At this point, 100µL-300µL of bacterial cultures (enough bacteria to yield a confluent layer in the soft agar in one day) was mixed with 3mL of 0.4% molten agar and poured onto the top of a KB agar plate. Top agar was allowed to solidify for ∼10 minutes, at which point a 10µL sample of each tailocin was added. Killing zones for each strain were observed and photographed either the next day or two days later depending on growth of the target strain. Overlay assays were carried out at least three independent times, with representative results reported. Unedited pictures of the overlay results can be found in supplemental files at doi: 10.6084/m9.figshare.22688020.

## Results

### Genomic Association for Tailocin Sensitivity Across P. syringae

There have been numerous switches in the phenotype of tailocin sensitivity across *P. syringae* strains [13]. We therefore hypothesized that genomic investigation of strains could highlight strong and independent signals pointing to the identity of genes and pathways contributing to differential tailocin sensitivity. The results of an analysis using Roary and Scoary to correlate genomic changes with tailocin sensitivity are shown in Table 2, and we find that 17 different loci are significantly (p<0.05) associated with tailocin sensitivity by these analyses and after Benjamini-Hochberg correction for multiple tests. A subset of 11 of these remain significantly associated with tailocin sensitivity after Bonferroni correction for multiple tests (p<0.05). Looking at the larger set more specifically, there are 9 loci whose presence is strongly correlated with tailocin sensitivity group A (as per [13]) and 8 whose presence is strongly correlated with tailocin sensitivity group B. Each of the 17 loci associated with each sensitivity group are colocated in syntenic genomic locations across strains, a location which corresponds to gene clusters implicated in production of the Common Polysaccharide Antigen (CPA) chain of the LPS as per [32].

To gain deeper insight into this genomic cluster and these results and to highlight diversification of this region between closely related strains, we focused on investigating gene presence and synteny within the entirety of the CPA locus across a subset of phylogroup 2 strains (Figure 1). We specifically found that this region was bracketed by conserved loci in nearly every strain, with the *ildD* coding for L-lactate dehydrogenase on one end and the gene for *ychF* coding for an ATPase on the other end. Genes found in this region in tailocin sensitivity group A include those involved in the production and modification of rhamnose moieties in the CPA (*gmd, rmd, wzm, algA,* and numerous glycosyl transferases and sugar modification enzymes). This region also encodes the TET operon, a type I secretion system to transport CPA sugar chains from the cytoplasm, but as highlighted in Jayaraman et al. [32, 33] and elaborated on here, there is extensive sequence variation between these operons across sensitivity groups. Genes found in this region in tailocin sensitivity group B also include those involved in the production and modification of rhamnose moieties in the CPA (*rfbABC,* AKA *rmlABC*) as well as other glycosyl transferases and sugar modification enzymes and two potential secretion systems). Despite diversity within this *ildD-ychF* region across these strains, it is clear that the genes implicated as contributing to tailocin sensitivity by the Scoary analysis are colocalized and found in syntenic locations across sensitivity classes. However, more pointedly, organization and sequence of this region completely changes with tailocin sensitivity and this change is consistent across all independent switches in sensitivity. Such changes appear to be much broader than that highlighted in [33, 34], as a much greater portion of this region is swapped out through what appears to be a recombination event.

**Figure 1.**
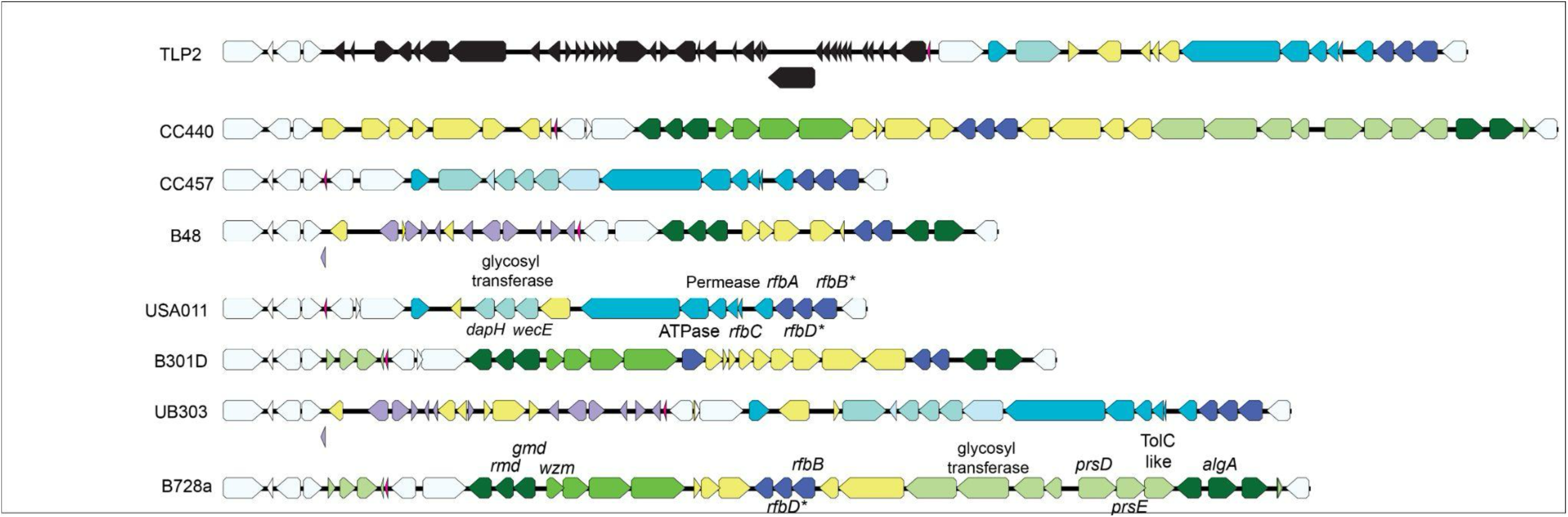
O-antigen / Common Polysaccharide Antigen Locus of *P. syringae* Phylogroup 2 Strains. The genomic island found between *ildD-ychF* is shown for 8 phylogroup 2 strains. Strains are arranged vertically in phylogenetic order according to Baltrus et al. 2019, and were picked for the figure because they represent situations where closely related strains have switched tailocin sensitivity class phenotypes. Common loci not thought to be involved in LPS construction are shown in white, and loci that are part of a possible prophage in strain TLP2 are shown in black. Loci that are possibly involved in LPS construction but which are found across tailocin sensitivity class strains are colored purple, with stronger coloration indicating presence in a larger number of strains. Yellow loci are only found in single strains among the eight shown. Loci that are possibly involved in LPS construction but which are found only within tailocin sensitivity class A and B strains are shown in green and blue, respectively, with stronger coloration indicating presence in a larger number of strains. Loci that were significantly associated with tailocin sensitivity class according to our analyses are labelled in strains *Psy*B728a (sensitivity class A) and USA011 (sensitivity class B), and we additionally label *rfbD* since we highlight its use as a predictor of tailocin sensitivity.

### rbfD As an Indicator Gene for Tailocin Sensitivity

A previous report identified gene sequences within the TET operon as fairly accurate indicators of tailocin sensitivity classes [33]. Upon inspection of the CPA locus across tailocin sensitivity classes, we observed that *rfbD* gene, encoding dTDP-4-dehydrorhamnose reductase, appears to be the only gene located in the *ildD-ychF* region across all strains. However, unlike the TET operon throughout *P. syringae*, sequence similarity across alleles of *rfbD* is extensive enough to be identified and analyzed across strains by sequence searches alone. We therefore pulled sequences for alleles of this locus from all strains from the main *P. syringae* phylogroups as described in [35] (phylogroups 1-6 and 10) and analyzed in [13], as well as 5 strains for which tailocin sensitivity has yet to be screened and inferred a maximum likelihood phylogeny from these sequences. We also include *rfbD* sequences from 11 strains for which LPS has been previously characterized but which also have yet to be screened for tailocin sensitivity. As one can see in Figure 2, alleles of RfbD form two distinct clades, with clade membership showing a complete match to tailocin sensitivity in all previously screened strains (with the caveat that tailocin sensitivity of three strains is misidentified in [13]; *Pto*T1, *Pla*106, CC1630 are labeled as sensitivity class A in this prior publication but should clearly be class B upon further comparison). From this phylogeny, we predict that strains *Pto*2170, Cit7, *Por* 1_36, *Phel* LMG 5067, and *Ptg* ICMP 4091 would be characterized as sensitivity class B,B,B,A and A respectively based on *rfbD* sequences. Furthermore, all strains where the LPS has been previously characterized [36] as dominated by D-rhamnose (*Pmp* CFBP 1650, *Ptg* ICMP 6370, *Phel* CFBP 1732) are predicted to be tailocin sensitivity class A while those dominated by L-rhamnose (*Pga* NCPPB 588, *Pga* NCPPB 1399, *Pga* NCPPB 2708, *Ppo* NCPPB 3545, *Pri* NCPPB 1010, *Pdl* NCPPB 1879) are predicted to be tailocin sensitivity class B. Lastly, two strains appear to maintain both L and D-rhamnose in their LPS based off of previous characterization (*Pta* NCPPB 79 and *Pla* NCPPB 1096), but their *rfbD* alleles clearly cluster with tailocin sensitivity class A.

**Figure 2.**
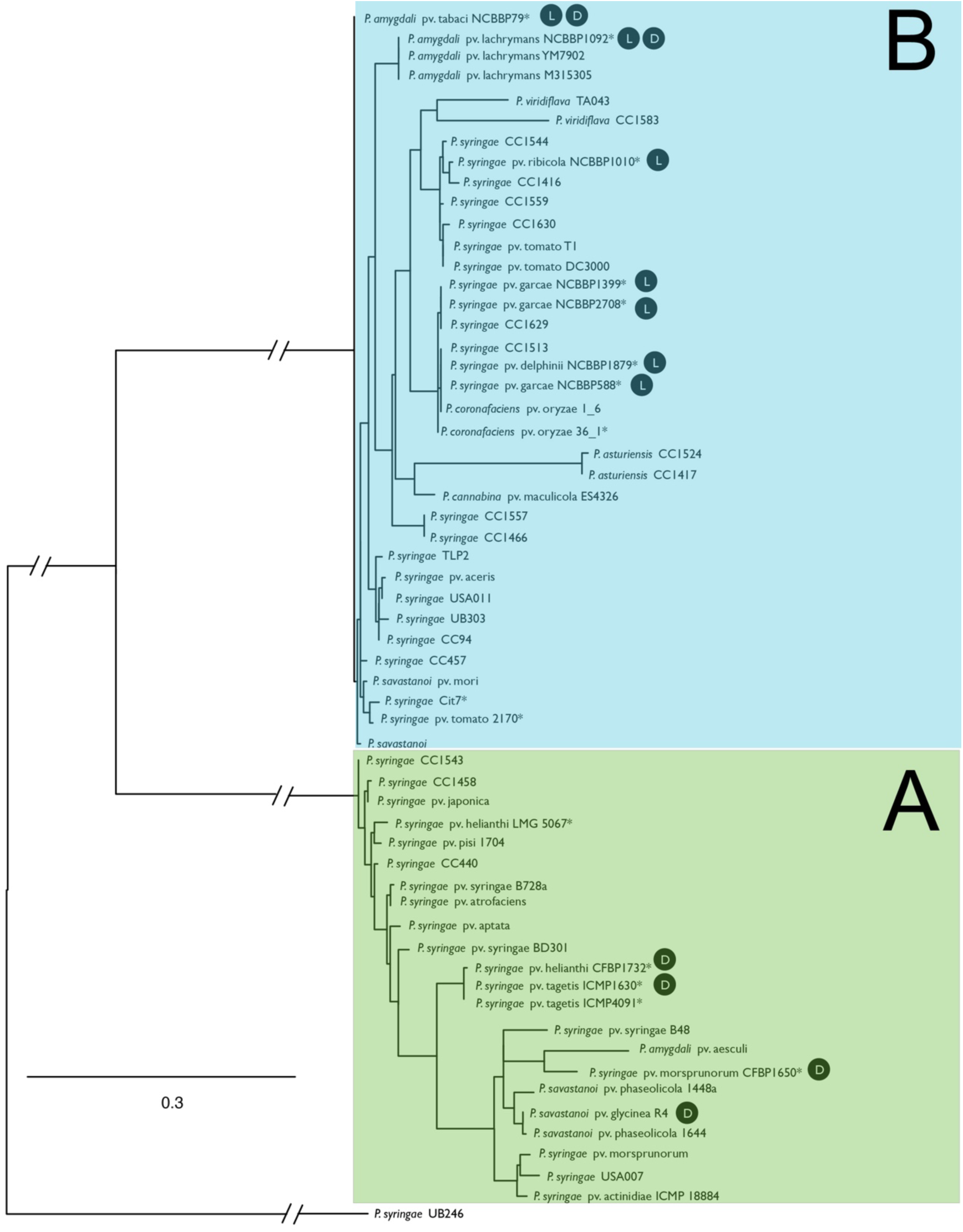
**RfbD as a Predictor of Tailocin Sensitivity Classes in *P. syringae.*** We inferred a maximum likelihood phylogeny from protein sequences of RfbD from across *P. syringae* strains tested for tailocin sensitivity in Baltrus et al. 2019 and from sixteen additional strains which have had their genomes sequenced but which have not been previously screened. Strain names are shown matching abbreviations in Table 1. Clades containing strains previously typed as tailocin sensitivity class A and B are colored green and blue, respectively. Alleles of RfbD from previously unscreened strains in the phylogeny, designate these strains with * and additionally, strains with characterized LPS conformations are noted with a black circle containing either “L” for L-rhamnose or “D” for D-rhamnose [36]. All files for phylogenetic analysis can be found in the supplemental data at doi: 10.6084/m9.figshare.22688020.

We performed overlay assays using these five strains which were not included in our previous tailocin sensitivity matrix and which represent multiple phylogroups of *P. syringae* as well as 11 strains for which LPS has been previously characterized (Figure 3). As a means to categorize tailocin sensitivity classes for these strains, we, used tailocins produced by four different strains from two phylogroups that delineate sensitivity groups A and B as per Baltrus et al. 2019: CC440 (phylogroup 2, killing class 1), USA011 (phylogroup 2, killing class 2), USA007 (phylogroup 3, killing class 1), and CC1416 (phylogroup 3, killing class 2). Results from these overlays are largely in line with predictions from *rfbD* phylogenies for strains Cit7, *Por* 1_36, and *Pto*2170, and show that these strains are classified as sensitivity group B (Figure 3B). When tailocin activity is witnessed, all strains where LPS has been previously shown to be dominated by D-rhamnose are tailocin class B while those dominated by L-rhamnose are tailocin class A with one exception (*Pga* NCCBP 588).

**Figure 3.**
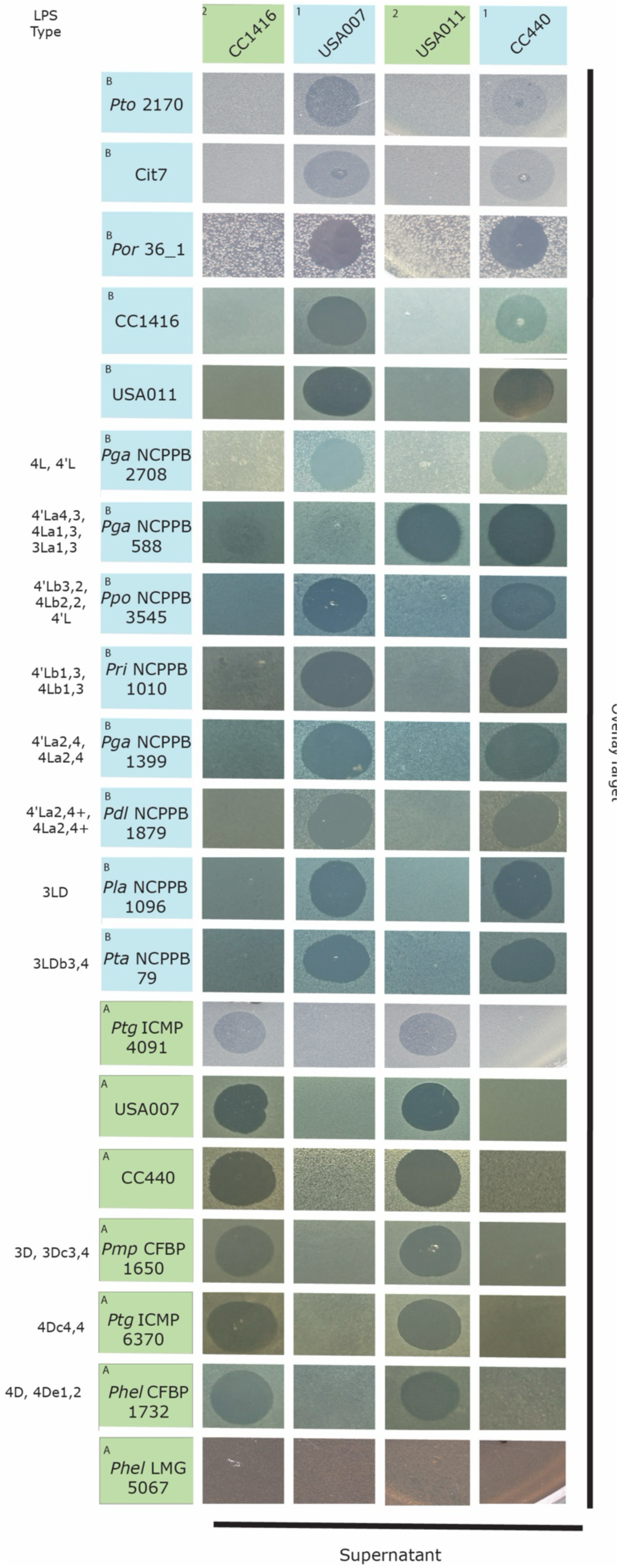
Tailocin Prediction for Unscreened Strains Largely Matches Predictions. We performed soft agar overlay assays where tailocins from four different strains and two different phylogenetic groups of *P. syringae* (USA007 and CC440, tailocin killing class 1, blue boxes; CC1416 and USA011, tailocin killing class 2, green boxes) were tested against sixteen strains for which tailocin sensitivity has not been previously reported as well as against themselves (strains listed in the Y axis, with strain names colored for predicted tailocin sensitivity class as per phylogeny in Figure 2A). Strain names are abbreviated as per Table 1. When strains being tested for sensitivity have had their LPS characterized, we list these characterizations to the left of their names with nomenclature shown according to [23].

### Numerous Predicted LPS Biosynthesis and Modification Genes Found on a Plasmid in Phylogroup 9 Strains

We also report two complete genome sequences for *P. syringae* phylogroup 9 strains CC1417 and CC1524, and highlight the presence of a plasmid conserved within both genomes. Notably, although this plasmid contains a handful of genes predicted to be associated with plasmid maintenance and replication (*repB,* a toxin-antitoxin system, *tus*) this replicon also contains numerous genes predicted to modify the LPS. These LPS-modification genes include the only predicted copy of *gmd* within the genome, a type I secretion system, and 8 different glycosyltransferases. This plasmid is highly conserved between both CC1417 and CC1524 and was circularized in each independent genome assembly. We also note that the main LPS biosynthesis region found of the chromosome of both strains, although containing genes like *rfbABCD*, is relatively reduced compared to all other strains reported within this manuscript.

## Discussion

Composition of the lipopolysaccharide layer of the outer membrane has been shown to impact multiple phenotypes crucial for survival of bacteria that are widely distributed throughout the environment [37]. Likewise, since the LPS often forms the interface between bacteria and hosts, changes in the LPS can have widespread effects during infection of plants by phytopathogenic bacteria [37, 38]. Our findings support previously published results implicating the LPS, and specifically a structure analogous to the CPA chain of *P. aerguinosa* formed by rhamnose polymers, as the target of *P. syringae* tailocins. Previous results suggested that changes in LPS composition across strains of pathovar *actinidae* were due to localized recombination of allelic variation of the TET operon [38], which is part of this CPA locus and encodes ABC transporter subunits, a bifunctional glycosyltransferase and an o-methyltransferase. The TET operon is responsible for the translocation, ligation, and modification of D-rhamnose moieties as they move from construction in the cytoplasm to the outer membrane. Jayaraman et al. were also able to extrapolate to strains outside of pathovar *actinidiae* and showed that allelic diversity in genes resembling the TET operon in context and sequence could predict tailocin sensitivity broadly across *P. syringae and P. viridiflava* strains.

While our results echo those of Jayaraman et al., the patterns reported here skew more towards presence/absence of genes and broad structural differences in the arrangement of the operons rather than *just* redistribution of allelic variation. Specifically, we show that operons encoding pathways for biosynthesis of precursors of the rhamnose moieties, multiple type I secretion systems including the TET operon, and various glycosyl transferases and sugar modification enzymes are syntenic and colocalized across many strains but also appear to undergo repeated wholesale swaps in gene content when a broader set of *P. syringae* genomes is analyzed for correlations with tailocin sensitivity. We further evaluate each of these operons in the context of LPS production below:

### gmd, rmd, algA

Alleles of *gmd* and *rmd* are differentially present within the CPA locus of strains classified as sensitivity class A (represented by strain *Pseudomonas syringae* pv. *syringae* B728a, *Psy*B728a, in Fig. 1) and form orthologous clusters according to the Roary pipeline which are significantly associated with tailocin sensitivity according to Scoary. Together with *wbpZ,* these genes are predicted to form an operon involved in the biosynthesis and linkage of D-rhamnose in the O-antigen of the LPS [39]. While spacing and orientation of these genes and the lack of factor independent transcriptional terminator suggests that all three could coregulated as an operon within *Psy*B728a, it is unclear whether regions upstream of *wbpZ* can independently control transcription of this gene.

D-rhamnose in the LPS is ultimately formed through the production of GDP-D-rhamnose from GDP-D-mannose. Gmd is predicted to encode GDP-mannose 4,6-dehydratase, an enzyme that carries out the first step in this pathway and converts GDP-D-mannose to GDP-4-dehydro-6-deoxy-D-mannose (aka GDP-6-deoxy-d-lyxo-hexos-4-ulose) [39]. Rmd is predicted to encode GDP-6-deoxy-d-lyxo-hexos-4-ulose-4-reductase, which produces GDP-D-rhamnose from the substrate produced by Gmd [39]. *wbpZ* is a glycosyltransferase that can attach D-rhamnose to the growing LPS chain from GDP-D-rhamnose [40]. Alleles of *algA* are encoded by genes within the CPA locus in strains from *P. syringae* tailocin sensitivity class A, also form an orthologous cluster according to the Roary pipeline, and are significantly associated with tailocin sensitivity class A strains according to Scoary. AlgA is predicted to encode a bifunctional protein with phosphomannose isomerase (PMI) and GDP-mannose pyrophosphorylase (GMP) activities, and which is therefore predicted to catalyze the conversion of fructose-6-phosphate to mannose-6-phosphate and mannose-1-phosphate to GDP-mannose [41]. It is likely that this enzyme catalyzes the substrate (GDP-mannose) that is acted upon by other genes in the CPA locus significantly associated with tailocin sensitivity in class A strains and which produce GDP-D-rhamnose, because GDP-mannose is the substrate for the Gmd protein. We note again that in contrast to strains from all other phylogroups, *gmd* appears to be plasmid bound in both phylogroup 9 strains (CC1417 and CC1524).

Many strains from *P. syringae* sensitivity class B can possess relatively distant orthologues of both *gmd* (∼80% protein identity) and *rmd* (∼30% protein identity). While they are positioned in close proximity within these strains, they are located outside of the CPA locus on the chromosome and do not appear to be closely associated with an additional glycosyltransferase. (data not shown). Disruption of this operon in sensitivity class A strain *P. syringae* pv. *actinidiae* ICMP 18884, and specifically within *gmd* (mutant B12), and *wbpZ* (mutant A1) has been shown to provide resistance to tailocin killing [44]. It has also been shown that a point mutation in *rmd* can provide tailocin resistance in the class A strain *P. syringae* pv. *phaseolicola Pph*1448a [42]. The significant association of D-rhamnose biosynthesis genes with tailocin sensitivity class A strains is strong evidence suggesting D-rhamnose as a potential binding target of tailocins that can kill these strains.

### rfbA, rfbB, wecE

Alleles of *rfbA* and *rfbB* from sensitivity class B form an orthologous cluster according to the Roary pipeline which are significantly associated with tailocin sensitivity according to Scoary. Additionally, an orthologue of *wecE* appears to be found within a separate operon within the CPA locus, but is also significantly correlated with tailocin sensitivity class B strains.

RfbA (also known as RmlA) is a glucose-1-phosphate thymidylyltransferase, which catalyzes the formation of dTDP-glucose from dTTP and glucose 1-phosphate [43]. It should be noted that there are two distinct enzymes (RfbA and RffH) identified in *E. coli* that can carry out this same reaction, and that these enzymes function in different pathways and are present in different operons in *E. coli* [44, 45]. RfbB is a dTDP-glucose 4,6-dehydratase, which catalyzes the dehydration of dTDP-D-glucose to form dTDP-6-deoxy-D-xylo-4-hexulose [46]. WecE is a dTDP-4-amino-4,6-dideoxygalactose transaminase [47]. L-rhamnose in the LPS is ultimately formed through the production of dTDP-L-rhamnose from GDP-L-mannose, and each of these enzymes is therefore predicted to be involved in the biosynthesis or modification of L-rhamnose for the O-antigen of tailocin sensitivity class B strains. As with the D-rhamnose biosynthesis gene associations with tailocin sensitivity class A genomes, the significant association of L-rhamnose biosynthesis genes with tailocin sensitivity class B strains strongly suggests L-rhamnose as a potential binding target of tailocins that can target these strains.

Lastly, we note that an independent orthologous cluster of *rfbB* alleles is also significantly associated with tailocin sensitivity class A strains. While this set of genes likely encodes an enzyme with similar functionality as those that are found in sensitivity class A strains, there is significant divergence between alleles of *rfbB* from each sensitivity class.

#### Potential Type I Secretion and Transporter Loci (*wzm,* ATPase, *prsD, prsE,* TolC-like, *rfbC,* Permease)

For class A genomes, the presence of an apparent orthologue of *wzm* as well an ATPase are significantly correlated with tailocin sensitivity. In *P. aeruginosa,* rhamnose precursors of the CPA are transported from the cytoplasm of the cell by an ABC transporter (*wzt/wzm*), with energy provided by an ATPase, and with a fourth glycosyltransferase gene present in the operon to attach these sugars to the growing chain [20, 48]. This particular operon in *P. syringae* is the same as highlighted by Jayaraman et al. for its patterns of recombination and it is likely that this operon in tailocin sensitivity class A strains is carrying out the same activities as the *wzt/wzm* ABC transporter in *P. aeruginosa*. Also worth noting is that there is a fourth gene encoding a glycosyltransferase (potentially referred to as *wbdD*) which appears to be part of this operon in sensitivity class A genomes. However, significant divergence for this gene within sensitivity classes due to recombination likely leads to a lack of overall significant associations overall for this enzyme with class A strains, as pointed out by Jayaraman et al. Disruption of this operon, and specifically within *wzm* (mutant A4)*, wzt* (mutant B4), and *wbdD* (mutants A6 and B6) has been shown to provide resistance to tailocin killing of the sensitivity class A strain *P. syringae* pv. *actinidiae* ICMP 18884 [50, 51]. Given the predicted enzymatic activities of these genes, it is likely that they are responsible for attaching D-rhamnose to the O-antigen of *P. syringae* tailocin sensitivity class A strains. This provides an additional line of evidence suggesting D-rhamnose as a potential binding target of tailocins that can target sensitivity class A strains.

Tailocin sensitivity class B genomes potentially encode multiple transporters that each contain subsets of genes significantly associated with tailocin sensitivity. One of these transporter operons appears to encode four genes that likely encode an ABC transporter used by Jayaraman in phylogenetic analyses as a predictor of tailocin sensitivity [52](and which is analogous to the potential *wzt/wzm* ABC transporter system mentioned above but divergent both in sequence and in annotation). Three of these genes in this operon are significantly associated with tailocin sensitivity for class B: a permease, *rfbC* (dTDP-4-dehydrorhamnose 3,5-epimerase), and an ATPase. As with the *wzt/wzm* operon mentioned above, there is often a fourth glycosyltransferase gene downstream of these three but which is divergent enough across strains so that significant associations with tailocin sensitivity are likely obscured. The presence of an *rfbC* orthologue as part of this pathway further hints that this may be the ABC transporter that enables building L-rhamnose chains as part of the *P. syringae* O-antigen, and provides additional support for the idea that L-rhamnose is a potential binding target of tailocins that can target sensitivity class B strains.

One additional type I secretion operon is significantly associated with tailocin sensitivity in many class B genomes, and contains genes similar to *prsD, prsE,* as well as a locus potentially encoding a TolC-like protein. Unlike the two transporter operons mentioned above, there does not appear to be a glycosyltransferase as part of this operon across strains. PrsD and PrsE are involved in exporting acidic polysaccharide in *Rhizobium* as part of biofilm formation, with PrsD acting as an ATP-binding protein and PrsE encoding a membrane fusion protein [49], but their functions within these *P. syringae* strains is unknown. TolC is an outer membrane channel protein for type I secretion systems [50]. By annotation, it is likely that this operon encodes proteins responsible for transporting sugars as part of the LPS of *P. syringae*, but at present it is difficult to identify the precise substrates. TolC outer membrane proteins are predicted to form “tunnels” through the outer membrane for transport of substrates for a variety of different type I secretion pathways.

#### Glycosyl-transferases

Sugar chains of the LPS are built through the action of glycosyltransferases, which take sugars linked to a variety of substrates and transfer them to the growing LPS chains [51, 52]. One glycosyltransferase, potentially found in an operon with *wecE* mentioned above, is significantly correlated with tailocin sensitivity group B strains and therefore is potentially involved in the transfer of L-rhamnose. One glycosyltransferase is significantly associated with tailocin sensitivity class A strains, and appears to be present as part of a larger operon across these strains, but is the only gene from this operon that shows a significant association. Outside of context clues, such as other enzymes encoded in operons with these loci, it is difficult to pinpoint the functions of these glycosyl transferases without further experiments. We also note that there appear to be multiple glycosyl transferases on plasmids of phylogroup 9 strains CC1417 and CC1524).

### dapH

*dapH* putatively encodes 2,3,4,5-tetrahydropyridine-2,6-dicarboxylate N-acetyltransferase, which catalyzes the transfer of an acetyl group from acetyl-CoA to tetrahydrodipicolinate [53]. There are multiple copies of similar enzymes throughout the *P. syringae* genome, but this particular copy is correlated with tailocin sensitivity group B genomes and potentially points towards the role of additional modifications of sugars in LPS production and tailocin sensitivity.

Previous manuscripts demonstrate that allelic variation within the TET operon could be effectively used for prediction of tailocin sensitivity [20]. Evaluation of genetic diversity in this pathway likely acts as a good metric for tailocin sensitivity when whole genome sequences are in hand, however, the TET operon shares little sequence similarity across strains within different tailocin sensitivity classes and thus there may not be an easy way to design probes to identify tailocin sensitivity classes without whole genomes. Our analyses further identify *rfbD* as potential additional candidate genes for extrapolating and predicting tailocin sensitivity classes from genome sequence alone for most *P. syringae* strains. Upon examination of the CPA locus of numerous *P. syringae* genomes, and unlike many of the genes significantly associated with tailocin sensitivity listed in Table 2, we observed that alleles of *rfbD* were always present within this island even though these alleles did not rise to the level of significance in the genome wide association study. Phylogenetic inference of RfbD clearly showed that sequences of each sensitivity group cluster with each other, and to the exclusion of sequences from the alternative sensitivity group, with strong support. Further, using this placement allowed for relatively high precision for assignment of previously unscreened strains to sensitivity classes. Strain *P. syringae* pv. *helianthii* LMG 5067 is predicted to be part of sensitivity class A by *rfbD* allele classification, but the lack of killing of this strain by either tailocin class likely points to biological nuances in LPS modification. Although RfbD appears critical for the creation of L-rhamnose [54, 55], a main component of the CPA chain of the LPS, it is unclear why there would be two different allelic clades for these genes across *P. syringae* classes and whether enzymes encoded by both clades have similar functionalities for the cell or whether each allele is optimized to work in concert with additional loci that are associated with each tailocin sensitivity class to encode LPS chains.

An abundance of data pulled together from different sources strongly suggests that the R type syringacins interact with rhamnose moieties that form the core chain of the O-antigen across *P. syringae* isolates. First, the R-type pyocins of *P. aeruginosa* have been shown to interact with rhamnose and glucose moieties in the LPS of target strains [9]. Although tailocins from *P. syringae* have been independently coopted from different phage than the R-pyocins, that these sugars have already been shown to be targeted by some tailoicns sets a prior precedent for evolutionary comparisons and predictions. More directly, studies characterizing the LPS of *P. syringae* demonstrate that the LPS of these strains fundamentally differs from what is known from *P. aeruginosa* and that strains can be classified into two LPS groups determined by whether the O-antigen is composed of repeating units of either D-rhamnose or L-rhamnose [36]. We report here that tailocin sensitivity differentially correlates with enzymes implicated in the synthesis of either D or L-rhamnose, with D-rhamnose chains implicated in sensitivity class A and L-rhamnose chains implicated for sensitivity class B and that these predictions are accurate for a subset of strains where the LPS has been previously characterized. Evidence from multiple strains also highlights that disruption of the production of D-rhamnose provides resistance to tailocins. Lastly, we report *rfbD* and tailocins classification for strains where the LPS has been previously characterized and show uniformly that strains containing only D-rhamnose LPS or L-rhamnose LPS are classified in tailocin sensitivity groups B and A, respectively. Thus, our predictions from genome-wide tailocin sensitivity associations do not only predict tailocin sensitivity based on *rfbD* alleles but also cleanly map onto known LPS conformations across strains. Taken together, our results create a compelling narrative that the chirality of rhamnose in the O-antigen of *P. syringae* at least partially determines classification for tailocin sensitivity: Chemical analysis suggests that *P. syringae* create either an L-rhamnose or D-rhamnose based O-antigen, genetic evidence shows that loss of O-antigen leads to tailocin resistance for most *P. syringae* tailocins where evaluated, and that two large scale tailocin sensitivity classes have been identified across *P. syringae* strains [13] and strongly correlate with presence/absence of operons involved in biosynthesis and transport of D-rhamnose and L-rhamnose. Assuming that this link does exist, it is worth noting that *P. syringae* pv. *glycinea* R4 is also categorized as tailocin sensitivity class A and possesses an O-antigen that is dominated by D-rhamnose moieties [56]. Future work will explore whether changing the chirality of the core CPA sugar is sufficient for changing tailocin sensitivity class. We also highlight that a recent publication build upon these results to demonstrate that structural variation in tail fibers of *P. syringae* tailocins correlate strongly with tailocin sensitivity and with *rfbD* alleles across strains, showing that “long” tail fibers associated with class 1 while “short” tail fibers of class 2 tailocins [57].

It is no surprise that composition of the LPS is extremely variable between bacterial isolates, as past studies have focused on using LPS characterization (in part) for classification of different lineages of *P. syringae* [36], but relatively few studies have evaluated the genomic basis of LPS diversity across *Pseudomonas syringae* (sensu lato). Our results here suggest that presence/absence diversity driven by relatively large-scale (∼20Kb) recombination events can dramatically and repeatedly alter LPS composition between closely related isolates and that such changes likely drive differences in sensitivity to different classes of tailocins. While the genetic basis of LPS configuration in *P. syringae* differs from canonical pathways established in *E. coli, P. aeruginosa* and other systems, in that the O-antigen appears to be dominated by enantiomers of rhamnose rather than a mix of variable sugar chains in addition to rhamnose, it is unknown how such variation impacts recognition of these strains by the plant immune system. It also remains unknown what correlated phenotypic changes, in the context of behaviors like biofilm formation or in responses to LPS interacting molecules like cationic antimicrobials [58], might occur across *P. syringae* strains with fundamentally different LPS conformations. It is also highly likely that phage attachment and infection also greatly contributes to, and may indeed itself drive, modification of the LPS under natural conditions [48, 59]. Our results highlight the potential for tailocins, or other LPS targeting bacteriocins [60], to be used as tools for uncovering and exploring genetic diversity in pathways and genes that underlie LPS production in these and other strains. Moreover, our study demonstrates that it is possible to uncover genetic and evolutionary patterns in LPS biosynthesis pathways by pairing large-scale phenotypic screens for tailocin sensitivity with genome sequencing, and such protocols could be used broadly to uncover interesting correlations in any system where strains display differential tailocin sensitivity.

There are multiple reports showing that independent recombination events occur at different locations in the chromosome in concert [13, 20], affecting LPS targeting ability and sensitivity across *P. syringae* strains. While there is currently no general mechanism identified to explain to how these independent recombination events co-occur, the coincident nature of these independent events hints that one of if not the main force driving fundamental changes in LPS composition within this phytopathogen over relevant evolutionary time scales is selection to prevent mismatches between tailocin killing capabilities and sensitivity. That these changes happen relatively frequently throughout the *P. syringae* phylogeny also suggests that the ability to diversity tailocin targeting capabilities, whether because of direct benefits during interstrain competition or in order to remedy phenotypic mismatches that may occur if alternative selection pressures shift LPS conformation prior to tailocin targeting, is an important trait for survival of natural *P. syringae* populations.

There are a two strains for which predictions from *rfbD* alleles don’t cleanly match observed tailocin sensitivity, *Ptg* ICMP 4091 and *Pga* NCCBP 588. In the case of *Ptg* ICMP 4091, phylogenetic relationships predict that this strain would be sensitivity class A while we observed that it was resistant to all four tailocins. This result is rare but not unprecedented and was seen with *P. syringae* pv. aceris in our previous manuscript [13]. *Pga* NCCBP 588 is predicted to be tailocin sensitivity group B, but is sensitive to supernatants produced by USA011 and CC440. In this case, we note that we used filtered supernatants rather than PEG precipitation for this assay, and that USA011 and CC440 are both classified as phylogroup 2 strains. Therefore, we believe that the killing activity witnessed is potentially not attributable tailocins, but rather due to an additional molecule produced by these strains in a phylogenetically conserved manner. Lastly, there are two strains for which LPS characterization has shown to maintain LPS with both L and D-rhamnose (*Pta* NCCBP 79 and *Pla* NCCBP 1096). In the assays presented we show that both strains are clearly categorized as sensitivity group B despite having both rhamnose moieties within the LPS. While not clearly shown in the three assay underlying data in Figure 3 (although slight inhibition can be seen), we note that we do occasionally see clear killing activity from both tailocin classes against these strains (data not shown). We therefore speculate that these strains can be killed by both tailocin types, but that LPS conformation within these strains is affected by additional environmental or growth conditions that leads to differential sensitivity.

Herein, we also report that at least two closely related phylogroup 9 strains (CC1417 and CC1524) maintain a unique configuration of LPS-biosynthesis and modification genes compared to all other *P. syringae* genomes. While all other strains referenced within this manuscript have at least a single dominant locus for LPS-biosynthesis found between *ildD* and *ychF*, these two phylgroup 9 strains also maintain a ∼100Kb secondary replicon that is highly biased towards genes predicted to impact LPS modification. Plasmid bound genes within these strains include *gmd* (coding for a critical enzyme for synthesis of D-rhamnose) as well as numerous glycosyltransferases. Although predicted LPS-modification genes can often be found in mobile elements such as plasmid and phage, this phylogroup 9 replicon is notable both for the proportion of predicted LPS-biosynthesis genes compared to other mobile elements within *P. syringae* as well as because to our knowledge *gmd* is found on the chromosome in all other strains. Although it is currently unclear how having such genes be plasmid-bound affects evolutionary and ecological dynamics of these strains with LPS-interacting particles (such as tailocins and phage), and these plasmids do not appear to contain genes for conjugation, this conformation highlights and emphasizes of how genomic plasticity can impact LPS-conformations. It is also notable, although potentially just a complete coincidence, that the phylogroup 9 strains are also the only *P. syringae* strains which appear to lack the genetic capacity to produce tailocins [13].

In sum, our results shed light on the potential genetic basis of fundamental switches in LPS conformation in the *P. syringae,* which appear to be strongly correlated with shifts in sensitivity to phage derived bacteriocions. They suggest that differences in binding of two previously identified tailocin killing types could in part be due to differential binding of rhamnose enantiomers that form the basis of the O-antigen of the LPS of this phytopathogen and point towards additional genomic loci that can be used to predict LPS and tailocin phenotypes from genomic sequences alone. Lastly, our study provides a clear proof of concept for combining large-scale phenotypic screens of tailocin sensitivity with genome wide association studies to identify the basis of bacteriocin interacting proteins and pathways.

## Notes

https://figshare.com/articles/dataset/Genomic_Correlates_of_Tailocin_Sensitivity/22688020

### Competing Interest Statement

The authors have declared no competing interest.

### Summary of Updates

We have included new experiments and genome sequences

